# *Bacillus licheniformis* global nitrogen homeostatic regulator TnrA is a direct repressor of *pgsBCAA* transcription in Poly-γ-glutamic acid synthesis

**DOI:** 10.1101/294728

**Authors:** Dongbo Cai, Yaozhong Chen, Shiyi Wang, Fei Mo, Xin Ma, Shouwen Chen

**Affiliations:** Environmental Microbial Technology Center of Hubei Province, Hubei Collaborative Innovation Center for Green Transformation of Bio-Resources, College of Life Sciences, Hubei University, Wuhan 430062, PR China

**Keywords:** *Bacillus licheniformis*, Poly-γ-glutamic acid, nitrogen regulator TnrA, transcription regulatory, γ-PGA synthase PgsBCAA

## Abstract

Poly-γ-glutamic acid (γ-PGA) is a multifunctional and naturally occurring biopolymer made from D- and L-glutamate as monomers, which is mainly produced by *Bacillus*. Few reports have been focused on the regulation network of γ-PGA synthesis in recent years. In this study, we have demonstrated that *Bacillus licheniformis* global nitrogen homeostatic regulator TnrA is a direct repressor of γ-PGA synthase PgsBCAA in γ-PGA synthesis. First, our results confirmed that TnrA repressed γ-PGA synthesis, deficiency of *tnrA* led to a 22.03% increase of γ-PGA production, and the γ-PGA yield was decreased by 19.02% in the TnrA overexpression strain. Transcriptional level assay illustrated that the γ-PGA synthase gene cluster *pgsBCAA* transcriptional level were increased in the *tnrA* deficient strain WXΔtnrA, indicating that γ-PGA synthase PgsBCAA was negatively regulated by TnrA. Furthermore, electrophoretic mobility shift assay (EMSA) and enzyme expression assays confirmed that TnrA directly repressed *pgsBCAA* expression by binding to *pgsBCAA* promoter, and the TnrA-binding site “CGTCGTCTTCTGTTACA” in the *pgsBCAA* promoter was identified by sequence and software analysis. Finally, computer analysis confirmed that the transcription regulations of γ-PGA synthase PgsBCAA by TnrA were highly conserved in other well-studied *Bacillus* species (*B*. *licheniformis*, *Bacillus subtilis* and *Bacillus amyloliquefaciens*). Collectively, our results implied that TnrA was a direct repressor for *pgsBCAA* expression in γ-PGA synthesis, and this research provided a novel regulatory mechanism underlying γ-PGA synthesis, and a new approach that deficiency of *tnrA* increases γ-PGA production.

**Importance:** γ-PGA is an important biopolymer with many applications, which is mainly produced by *Bacillus* species. Glutamic acid is the precursor for γ-PGA synthesis, which is catalyzed by the γ-PGA synthase PgsBCAA. Previously, the expression of PgsBCAA was reported to be regulated by ComA-ComP and DegS-DegU, DegQ and SwrA systems, however, few researches were focused on the regulation network of γ-PGA synthesis in recent years. In our research, the γ-PGA synthase PgsBCAA was confirmed to be negatively regulated by the nitrogen metabolism regulator TnrA, and the TnrA binding site in the *pgsBCAA* promoter was identified in *B. licheniformis* WX-02. Furthermore, computer analysis implied that TnrA-mediated regulation effect on *pgsBCAA* expression was highly conserved in *Bacillus*. Collectively, our research provided a novel regulatory mechanism underlying γ-PGA synthesis, and a new approach that deficiency of *tnrA* increases γ-PGA production.

## Introduction

Poly-γ-glutamic acid (γ-PGA) is a multifunctional and naturally occurring biopolymer made from D- and L-glutamate as monomers (1). Due to its owns the features of cation chelating, hygroscopicity, water-solubility, non-toxicity and biodegradability, γ-PGA has been used as various materials such as drug carriers, food preservatives, metal ion chelating agents, highly water absorbable hydrogels and fertilizer accelerators, and applied in the areas of medicine, food, water treatment, cosmetics and agriculture etc (2, 3).

Generally, *Bacillus* species were regarded as the efficient γ-PGA producers, and the precursor, *in vivo* glutamic acid, was applied for γ-PGA biosynthesis under the catalysis of γ-PGA synthase complex PgsBCAA (2). Among this gene cluster, gene *pgsB* acts as the main function role of γ-PGA synthase, whereas *pgsC* owns the function of γ-PGA secretion, and *pgsAA* is necessary for the next monomer addition and transportation of γ-PGA through the cell membrane (4). Previously, the γ-PGA synthesis was reported to be regulated by the two-component systems ComA-ComP and DegS-DegU, DegQ and SwrA systems (5). Among them, ComA-ComP and DegS-DegU, DegQ were involved in the quorum sensing, osmolarity, phase variation signals, while SwrA appears to act at the post-transcriptional level. Based on the previous researches, the regulation of γ-PGA synthesis by DegQ was well investigated, and the γ-PGA synthesis capability was abolished in the *degQ* mutant strain. Phosphorylated DegU (DegU-P) and SwrA up-regulated the expression level of *pgs* operon for γ-PGA synthesis (6, 7), and no more in-depth study was conducted to resolve this mechanism. Recent years, several researches were focused on the improvement of γ-PGA production by metabolic engineering strategies (8–10), however, few works were conducted to elaborate the regulation network of γ-PGA synthesis, thereby, it was unclear that whether the γ-PGA synthase PgsBCAA was regulated by other regulators.

The global nitrogen homeostatic regulator TnrA is known as a member of MerR family transcriptional factor, and it could both activate and repress the expression of many genes under nitrogen limited condition (11). Generally, TnrA up-regulated the expression levels of genes encoding ammonium transport (*amtB*-*glnK*) (12), nitrate assimilation (*nasAB*) (13), nitrite assimilatory enzymes (*nasDEF*) etc (14), and also exerted negative effects on the expression of *glnRA* (encoding for glutamine synthase) (15), *gltAB* (encoding for glutamic acid synthase) (16), *ilv*-*leu* (encoding for branched-chain amino acids synthase) (17), *degU* (encoding for two-component system DegS-DegU) etc (18). Previously, the chromatin immunoprecipitation coupled with hybridization to DNA tiling arrays (ChIP-on-chip) was applied to identify the TnrA binding sites at the genome scale of *B. subtilis*, and their results provided that TnrA binds reproducibly to 42 regions. Among them, 35 TnrA primary regulons were confirmed by real time in vivo transcriptional profiling using firefly luciferase. In their research, a predicted TnrA box has been implied in the promoter region of γ-PGA synthase gene *pgsB* (19), however, no in-depth research was conducted to analyze the relationship between TnrA and *pgsBCAA* expression.

*Bacillus licheniformis* WX-02 was proven as an efficient γ-PGA producer (20), and our previous research found that the genes related to glutamic acid synthesis were activated by 6% NaCl addition (21), and the glutamic acid synthesis capabilities were also improved by the physicochemical stresses such as heat, osmotic and alkaline etc (22–24). Furthermore, several metabolic engineering strategies have been conducted to improve the γ-PGA production (8, 25, 26). In this study, we demonstrated that *Bacillus licheniformis* global nitrogen homeostatic regulator TnrA is a direct repressor of *pgsBCAA* transcription in γ-PGA synthesis, and the TnrA-mediated regulation effect on *pgsBCAA* expression was highly conserved in *Bacillus*.

## Results

### Deficiency of *tnrA* increased γ-PGA production

TnrA is known as a global nitrogen regulator in *B. subtilis*. In order to test the function role of TnrA on γ-PGA production in *B. licheniformis*, the *tnrA* deficient and overexpression strains were constructed based on *B. licheniformis* WX-02, and the recombinant strains were named as WXΔtnrA andWX/pHY-tnrA, respectively. Then, these recombinants, as well as the control strains WX-02 and WX/pHY300, were cultivated in the γ-PGA production medium, and the γ-PGA yields at the end of fermentation (32 h) were measured to evaluate the effects of *tnrA* deficiency and overexpression on the γ-PGA production. Based on our results of **Fig. 1**, WXΔtnrA produced 38.37 g/L γ-PGA, increased by 22.03% compared with that of WX-02 (31.45 g/L), indicating that deficiency of *tnrA* improved γ-PGA production. Meanwhile, overexpression of TnrA led to a 19.02% decrease of γ-PGA yield, compared to WX/pHY300.

**Fig. 1:**
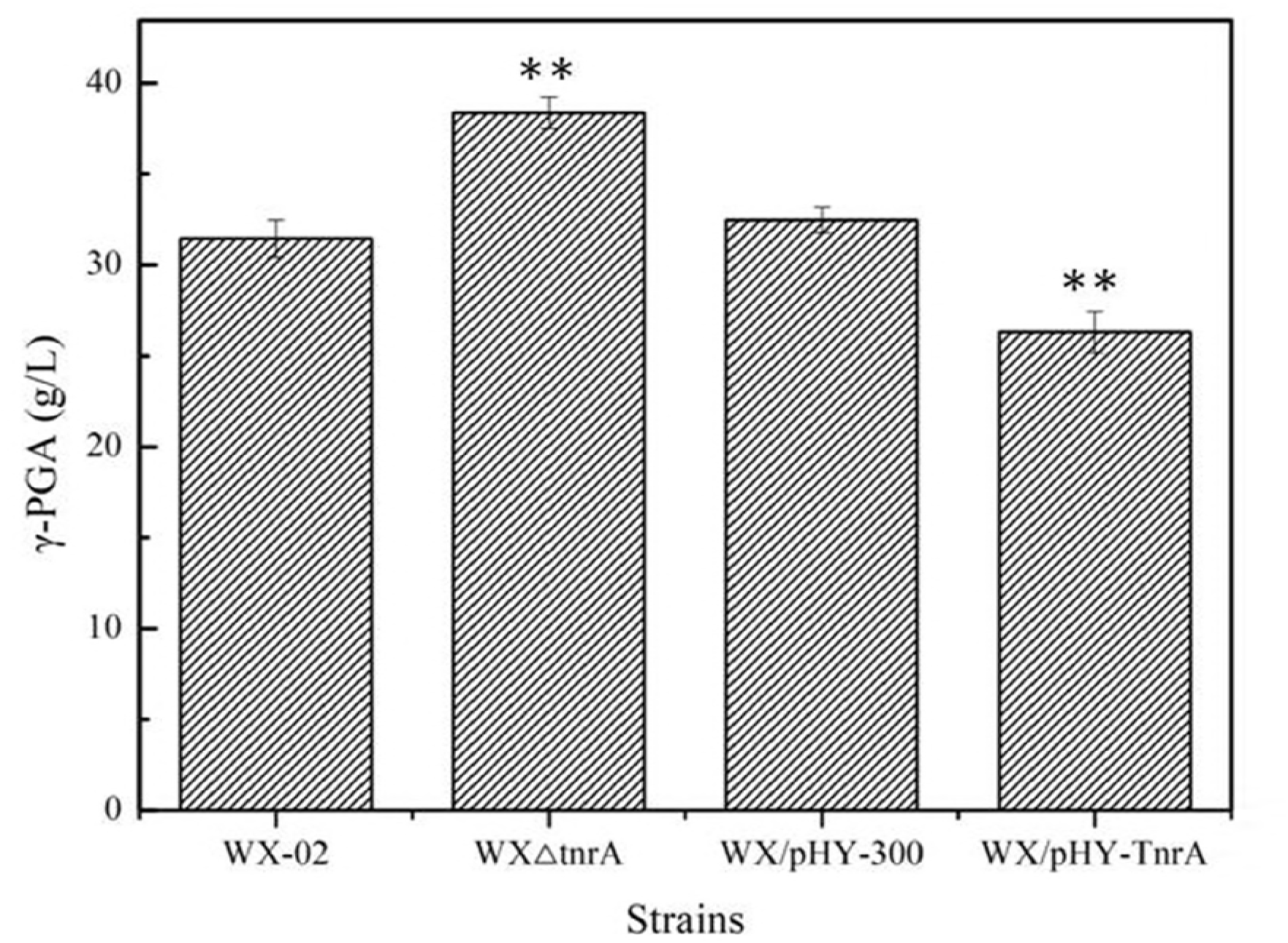
Effects of *tnrA* deficiency and overexpression on γ-PGA production. Data are represented as the means of three replicates and bars represent the standard deviations, ∗, P < 0.05; and ∗∗, P < 0.01 indicate the significance levels between recombinant strains and control strain.

Furthermore, the γ-PGA yields, cell biomass, residue concentrations of glucose and glutamic acid of these strains were measured during γ-PGA production. Our results implied that the γ-PGA yields of WXΔtnrA were higher than those of WX-02 throughout the whole fermentation process, and the maximum γ-PGA yield was increased by 22.03%. Glucose and glutamic acid consumption rates of WXΔtnrA were 9.13% and 11.34% higher than those of WX-02, respectively. Additionally, since the deficiency of *tnrA* affects the expression levels of nitrogen utilization genes (19, 27), which led to the low cell growth of WXΔtnrA during the γ-PGA fermentation. The specific γ-PGA yield of WXΔtnrA was 6.11g/g _DCW_, increased by 42% compared with that of WX-02 (4.31g/g _DCW_). Meanwhile, overexpression of *tnrA* decreased the γ-PGA production, as well as the glucose and glutamic acid consumption rates, and the cell biomass increased obviously in the *tnrA* overexpression strain (**Fig. 2**). Collectively, our results demonstrated that the nitrogen regulator TnrA repressed the synthesis of γ-PGA, and deficiency of *tnrA* increased γ-PGA yield.

**Fig. 2:**
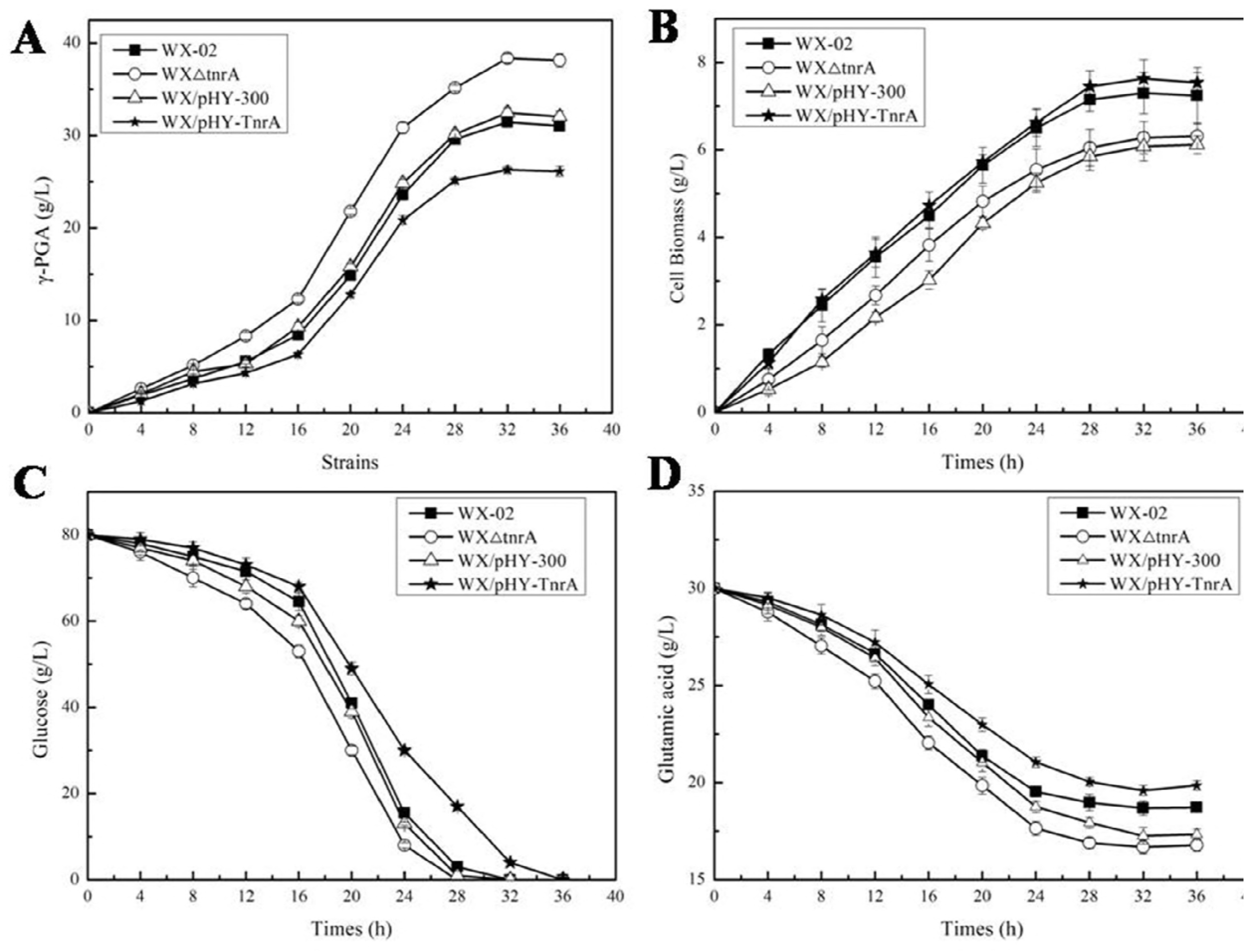
Fermentation process curve of *B. licheniformis* WX-02, WXΔ tnrA, WX/pHY300 and WX/pHY-TnrA. **A**: γ-PGA; **B**: Cell biomass; **C**: Glucose concentrations; **D**: Glutamic acid concentrations.

### The transcriptional levels of γ-PGA synthase genes *pgsBCAA* were increased in the *tnrA* deficient strain

Then, the transcriptional level of genes involved in glucose metabolism and γ-PGA biosynthesis were determined during γ-PGA production. As shown in **Fig. 3**, no *tnrA* transcriptional level was determined in WXΔtnrA, indicating that *tnrA* was deleted successfully. The transcriptional levels of γ-PGA synthesis genes *pgsB*, *pgsC* and *pgsAA* were all increased in WXΔtnrA (5.53-fold, 4.64-fold and 4.08-fold, respectively), and the transcription levels of their regulator genes, *degU*, was increased by 1.68-fold in the *tnrA* deficient strain. The glutamate synthase gene *gltA* was increased by 2.56-fold in WXΔtnrA. Moreover, the transcriptional levels of glucose-6-phosphate isomerase gene *pgi* and glyceraldehyde-3-phosphate dehydrogenase gene *gapA* in the glycolytic pathway were increased obviously, as well as the citrate synthase gene *citB* and isocitrate dehydrogenase gene *icd* in tricarboxylic acid (TCA) cycle. The glucose 6-phosphate dehydrogenase gene *zwf* in pentose phosphate pathway showed no significant changes in *tnrA* deficient strain. Furthermore, the genes, *alsS* and *alsD*, which responsible for the synthesis of main byproducts acetoin and 2,3-butandieol, were all decreased in WXΔtnrA, indicating that the deficiency of *tnrA* repressed the overflow metabolism, which were beneficial for γ-PGA synthesis.

**Fig. 3:**
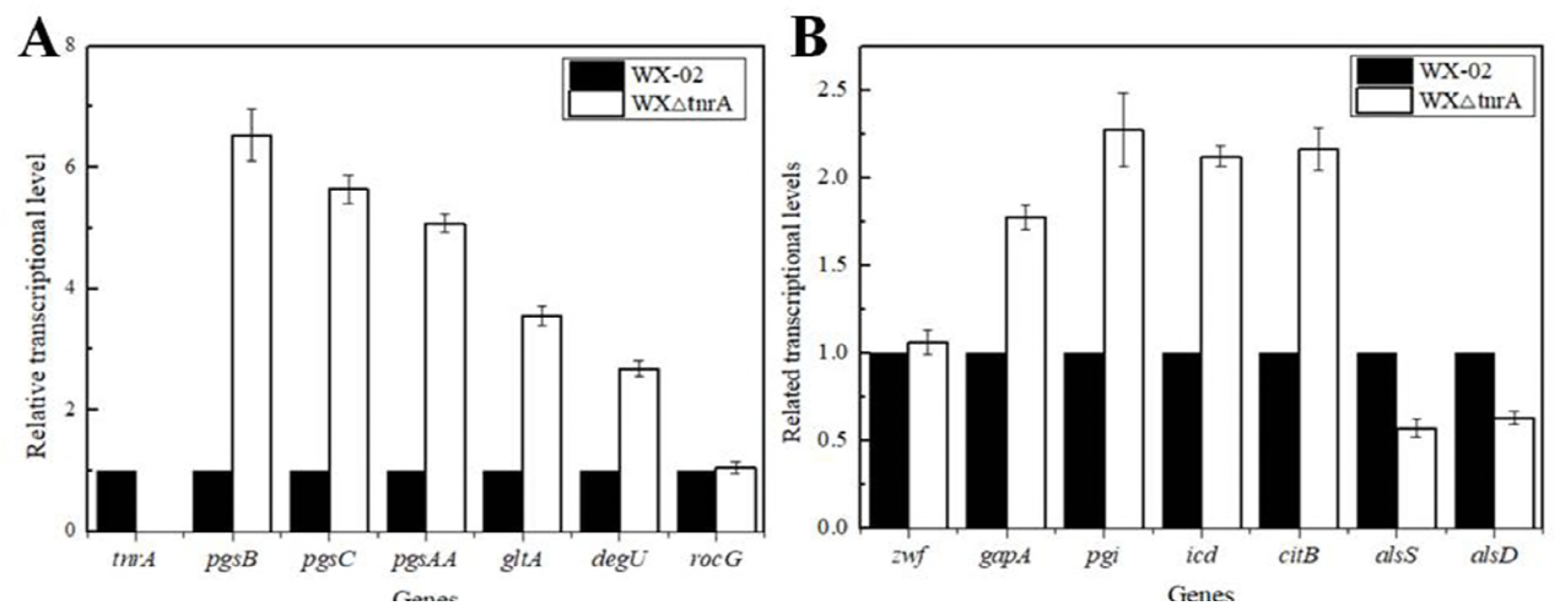
Transcriptional level analysis. **A:** Effects of *tnrA* deficiency on the relative transcriptional levels of genes in γ-PGA biosynthesis; **B:** Effects of *tnrA* deficiency on the relative transcriptional levels of genes in glucose metabolism.

### Identification of TnrA binding sites in the *pgsBCAA* promoter

Previous research has implied that there might be a predicted TnrA box in the *pgsB* promoter of *B. subtilis* (19). To verify whether TnrA regulated the *pgsBCAA* promoter directly or not in *B. licheniformis* WX-02, electrophoretic mobility shift assays (EMSAs) were performed. We expressed His_6_-tagged TnrA in *E. coli* BL21(DE3) and purified, and the SDS-PAGE results in **Fig. 4A** demonstrated that the TnrA protein was induced and purified successfully. Then, the DNA probes of *pgsBCAA* promoter (form -300 to +50) were amplified, and the purified DNA probes were incubated with the purified His_6_-tagged TnrA according to the instruction manual of EMSA kit. As shown in **Fig. 4B**, a TnrA-P_*pgsB*_ complex was formed in a concentration-dependent manner, indicating that TnrA could bind to the promoter of *pgsBCAA*.

**Fig. 4:**
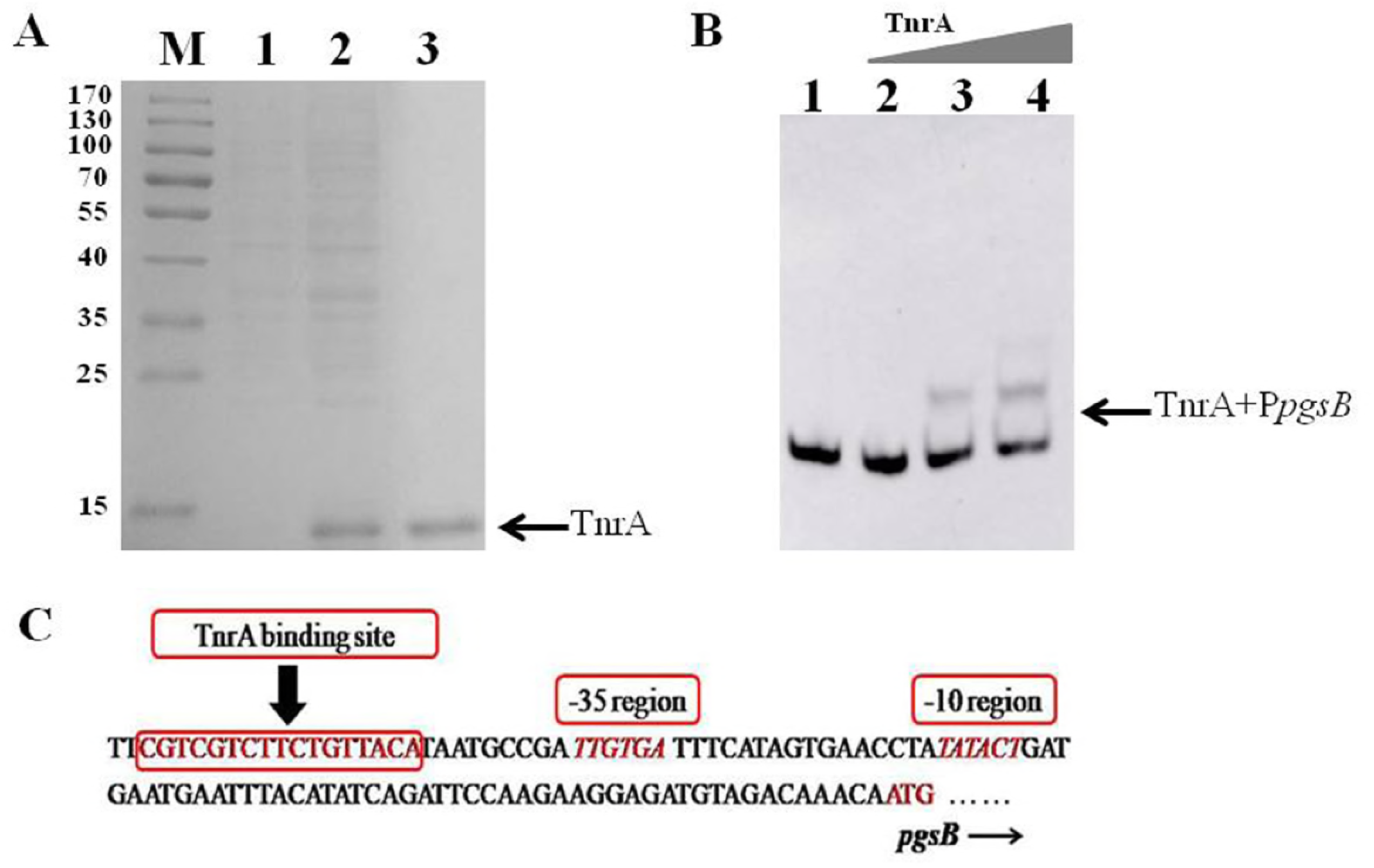
EMSA assay and identification of the TnrA binding site in the *pgsBCAA* promoter. **A: Expression and purification of TnrA in BL21(DE3);** Lane 1: The protein Marker (170 DKa, 130 DKa, 100 DKa, 70 DKa, 55 DKa, 40 DKa, 35 DKa, 25 DKa, 15 DKa,), Lane 2: The total intracellular protein of BL21/pET-TnrA before induction, Lane 3: The total intracellular protein of BL21/pET-TnrA induced by IPTG, Lane 4: The purified TnrA protein. The arrow indicates the purified TnrA (13.17 KDa); **B: EMSA assay of the purified TnrA protein and *pgsBCAA* promoter.** Lane 1: EMSA of TnrA protein (200 ng) with biotin-labeled *pgsBCAA* promoter P_*pgsB*’_, which lacking out of the TnrA binding site “CGTCGTCTTCTGTTACA”, as the negative group; Lane 2-4: EMSA of different TnrA concentrations (0, 100 ng, 200 ng) with biotin-labeled *pgsBCAA* promoter (-300 to +50 bp upstream and downstream of the translational start), and the arrow indicates the mixture of TnrA with *pgsB* promoter. **C: Identification of the TnrA-binding site in the *pgsBCAA* promoter of *B. licheniformis* WX-02**.

Previously, the TnrA box was proven to be a 17-bp interrupted, inverted repeat sequence “TGTNANAWWWTNTNACA” in *B. subtilis* (28). In this research, based on the conserved sequence of TnrA box and MEME software screening, the suspected TnrA binding site in *pgsBCAA* promoter, “CGTCGTCTTCTGTTACA”, was screened, which is outside of the “-10” and “-35” regions of *pgsBCAA* promoter (**Fig. 4C**). Then, the sequence of predicted TnrA box was further deleted in the *pgsBCAA* promoter, and formed the promoter P_*pgsB*’_. Our results confirmed that there was no TnrA-P_*pgsB*’_ complex formed in the EMSA assay, even though 200 ng TnrA was applied **(Fig. 4B)**. Thus, it was confirmed that TnrA directly band the *pgsBCAA* promoter, and the TnrA box “CGTCGTCTTCTGTTACA” in *pgsBCAA* promoter has been identified in *B. licheniformis* WX-02.

### *pgsBCAA* promoter is negatively regulated by TnrA

Furthermore, to investigate whether *pgsBCAA* was negatively or positively regulated by TnrA *in vivo*, the nattokinase expression assay was performed. Firstly, to exclude the interference of regulator DegU, which has been confirmed as the direct positive activator of *pgsBCAA* in the previous research (6, 7), we deleted gene *degU*in WX-02 and WXΔtnrA, and resulting in the mutant strains WXΔdegU and WXΔtnrAΔdegU. Then, the nattokinase expression vector mediated by P_*pgsB*_ were constructed, and electro-transferred into WXΔdegU and WXΔtnrAΔdegU, respectively. Based on our results of **Fig. 5**, the nattokinase activities of WXΔtnrAΔdegU were higher those of WXΔdegU under the condition of P_*pgsB*_ promoter, and the maxmium activity of WXΔtnrAΔdegU was 26.39 FU/mL, increased by 45.19% compared with that of WXΔdegU (18.18 FU/mL). Also, the specific nattokinase activity of WXΔtnrAΔdegU/pPgsBSacCNK was 2.57 FU/OD_600_, increased by 91.66% compared to WXΔdegU/pPgsBSacCNK (1.34 FU/OD_600_) (**Fig. 5A**). Thus, these results indicated that deficiency of *tnrA* might improve the expression of *pgsBCAA* promoter, which led to the improvement of nattokinase activity in the WXΔtnrAΔdegU/pPgsBSacCNK. Additionally, to exclude the interference of *tnrA* deficiency on the cell growth, which might affect nattokinase production, the nattokinase vector mediated by P43 prmoter was applied, and electro-transferred into WXΔdegU and WXΔtnrAΔdegU, respectively. Our results implied that the nattokinase activities produced by WXΔtnrAΔdegU/pP43SacCNK were lower than those of WXΔdegU/pP43SacCNK throughout the fermentation process (**Fig. 5B**), and this might due to that deletion of *tnrA* led to the reduction of cell growth, which further influence nattokinase production (27). Collectively, these above results illustrated that TnrA negatively regulated the transcription of *pgsBCAA*, and the nattokinase activity mediated by P_*pgsB*_ was improved in the *tnrA* deficient strain.

**Fig. 5:**
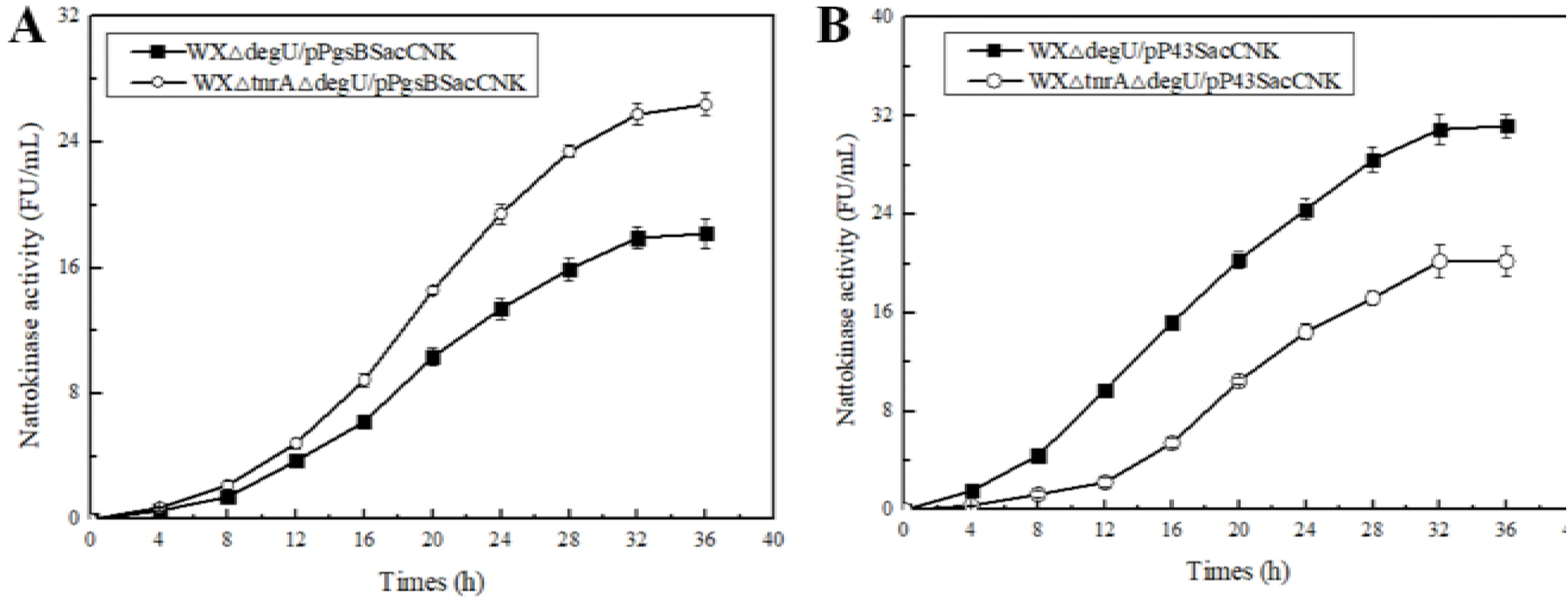
Confirmation of the regulation model of γ-PGA biosynthesis by TnrA via nattokinase expression assay. **A:** nattokinase activity produced by WXΔdegU/pPgsBSacCNK and WXΔtnrAΔdegU/pPgsBSacCNK; **B:** nattokinase activity produced by WXΔdegU/pP43SacCNK and WXΔtnrAΔdegU/pP43SacCNK.

### The transcriptional regulation of *pgsBCAA* by TnrA is highly conserved in *Bacillus*

Finally, we investigated whether TnrA directly regulate the transcription of γ-PGA synthase PgsBCAA or not in other *Bacillus* species. We screened the γ-PGA synthase gene cluster *pgsBCAA* in three well-studied *Bacillus* species (*B. licheniformis*, *B. subtilis*, *B. amyloliquefaciens*) according to the genome annotation from NCBI database, and searched the TnrA binding motif in their promoters regions. Sequence and software analyses indicated that TnrA binding sites were located in the *pgsBCAA* promoters of various *Bacillus* species: 14/15 *B. licheniformis*, 75/76 *B. subtilis*, 21/22 *B. amyloliquefaciens*. Meanwhile, the TnrA boxes were predicted in the corresponding promoters, which all located outside of the “-10” and “-35” regions of *pgsBCAA* promoters (**Table 3**). Thus, these results suggested that γ-PGA synthase PgsBCAA regulated by nitrogen regulator TnrA was highly conserved in *Bacillus*.

## Discussion

γ-PGA is an important biopolymer with many applications (29). Recently, several researches have been focused on the strategies of metabolic engineering to improve γ-PGA production, including eliminating the byproduct synthesis pathways, strengthening the γ-PGA pathways, manipulating ATP supply and NADPH generation (8, 10, 25, 30). However, the regulation network of γ-PGA synthesis has not been well analyzed, besides the regulators ComA~ComP, DegS~U, DegQ and SwrA (**Fig. 6**). In this study, it was confirmed that the nitrogen regulator TnrA was a directly repressor of γ-PGA synthase PgsBCAA, and deficiency of *tnrA* led to a 22.03% increase of γ-PGA yield.

**Fig. 6:**
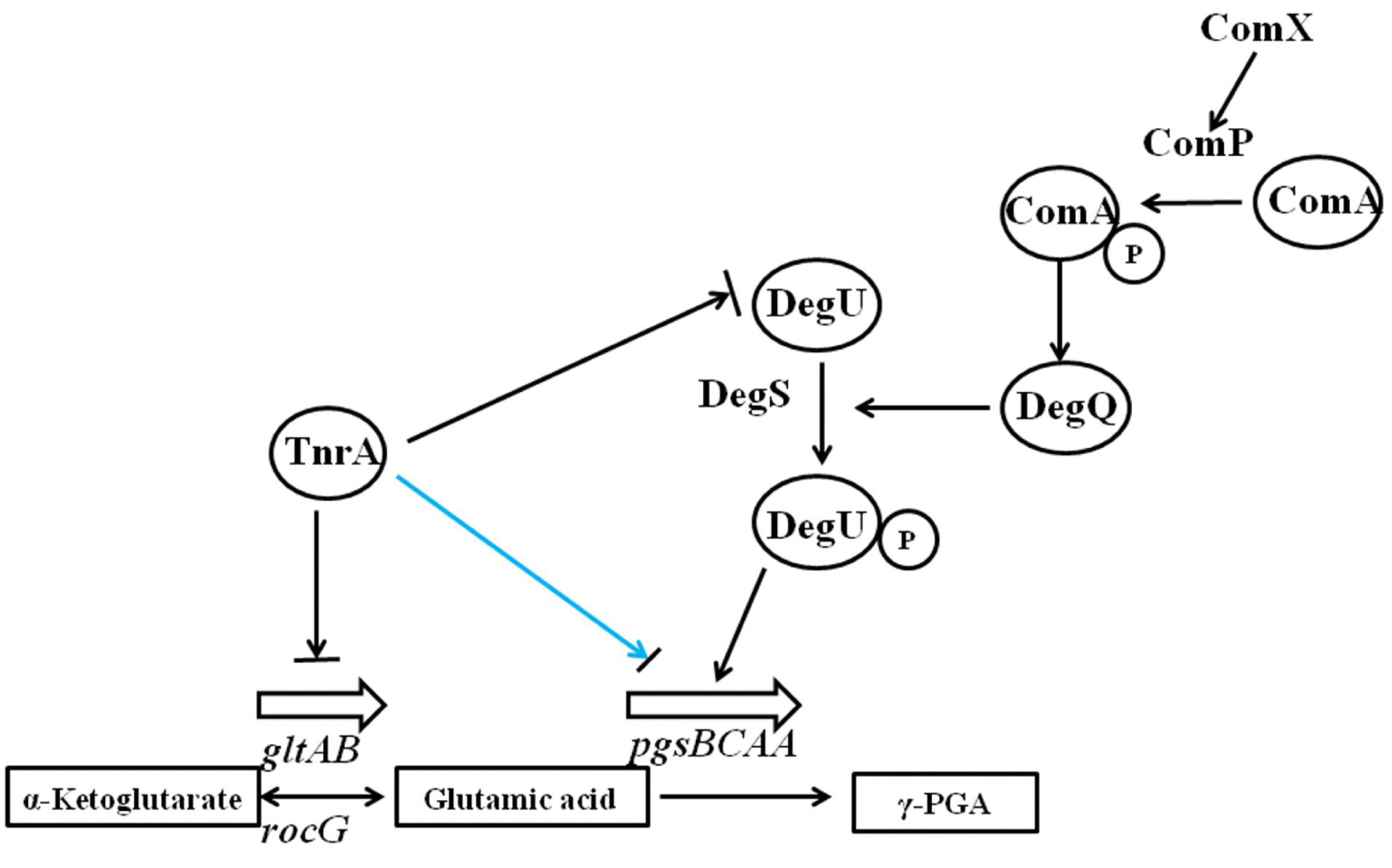
The proposed regulatory network of TnrA in γ-PGA biosynthesis in *B. licheniformis*. (The black lines indicate the previously proven regulatory mechanisms in *Bacillus*, the cyan line indicates the regulatory mechanism confirmed in this research).

γ-PGA was mainly produced by *Bacillus* species. *In vivo* glutamic acid is served as the precursor for γ-PGA synthesis, and nitrate addition could strengthen γ-PGA synthesis by our previous research, and ammonium salt was also acted as the important resource for γ-PGA synthesis (24, 31). Thus, nitrogen metabolism played an important role in γ-PGA synthesis. Since TnrA is a global nitrogen metabolism regulator in *Bacillus*, which acted both as the activator or repressor for nitrogen metabolism genes (11), thus, the regulator TnrA might also affect γ-PGA synthesis. Based on the previous research, a suspected TnrA box was predicted in the *pgsB* promoter in *B. subtilis* by Chip-on-chip (19), and our research confirmed that the γ-PGA synthase complex PgsBCAA were directly regulated by the nitrogen regulator TnrA, and the related genes (*pgsB*, *pgsC* and *pgsAA*) transcriptional levels were all increased obviously in the *tnrA* deficient strain, which led to the improvement of γ-PGA yield. In addition, TnrA was also proven as the repressor of the genes *gltAB* and *degU* (16), which were responsible for glutamate synthase and regulation of γ-PGA synthase in *Bacillus*. Based on our results, *gltA* and *degU* transcriptional level were increased obviously in the *tnrA* deficient strain, and this might be another reason that the γ-PGA yield produced by WXΔtnrA was higher than that of WX-02, and the proposed regulatory network of TnrA in γ-PGA biosynthesis in *B. licheniformis* WX-02 was provided in **Fig. 6**. Additionally, the nitrogen regulators GlnR and CodY acted as the similar features compared to TnrA, however, whether these regulators affect γ-PGA synthesis or not is still unknown recent now.

Previously, the γ-PGA synthesis was reported to be regulated by the two-component systems ComA-ComP and DegS-DegU, DegQ and SwrA systems (5). The γ-PGA synthesis capability was abolished in the *degQ* mutant strain, and phosphorylated DegU (DegU-P) and SwrA could up-regulate the expression level of *pgs* operon for γ-PGA synthesis (6, 7). Since TnrA was reported to be the repressor for *degU* expression (18), the transcriptional level of *degU* was increased by 1.68-fold in the *tnrA* deficient strain, thus, TnrA could affect γ-PGA synthesis through up-regulating the expression levels of *degU*, and the increase of *degU* transcriptional level was beneficial for γ-PGA synthesis. In our work, in order to exclude the interference of regulator DegU, we deleted gene *degU* in WX-02 and WXΔtnrA, and further analyze the regulation mechanism of *pgsBCAA* promote by TnrA in the *degU* mutant strains. Our results confirmed that nattokinase activity produced by WXΔtnrAΔdegU/pPgsBSacCNK was increased obviously compared to WXΔdegU/pPgsBSacCNK. Additionally, the TnrA-binding site “CGTCGTCTTCTGTTACA” was identified in the *pgsBCAA* promoter of *B. licheniformis* WX-02. Therefore, our results confirmed that TnrA could directly repress the transcription of *pgsBCAA* by binding to its promoter, in addition to the indirect effect of DegU. Nonetheless, the in-depth research is worth to explore that which pathway plays the major role for the improvement of γ-PGA yield in the *tnrA* mutant strain.

*Bacillus* species were the important industrial production strains (32), and they could produce various kinds of bio-chemical products (33), also, *Bacillus* could be acted as the host strains for protein expression (34). Among them, *B. subtilis*, *B. licheniformis*, *B. amyloliquefaciens* are the three well-studied *Bacillus* species, and they are also the main producers for γ-PGA synthesis (29). Furthermore, these species have a high degree of homology, *B. licheniformis* owns a nearly 75% identity with *B. subtilis* at the genome-level, and *B. amyloliquefaciens* has been regarded as a subspecies of *B. subtilis*, which genome identity is more than 80% with *B. subtilis*. Thus, these species might belong to the same branch in evolutionary terms. Based on our results, the TnrA boxes of *pgsBCAA* promoters were predicted in the most of these strains (**Table 3**), thus, our results suggested that the regulation of γ-PGA synthase PgsBCAA by TnrA might be conserved in *Bacillus*.

In conclusion, the regulation model of γ-PGA synthase PgsBCAA by nitrogen regulator TnrA was analyzed in this study. Based on our results, the γ-PGA synthesis was negatively regulated by TnrA, and deficiency of *tnrA* led to a 22.03% increase of γ-PGA yield. Transcriptional analysis confirmed that *pgsBCAA* expression level was all increased in the *tnrA* deficient strain, and EMSA and enzyme expression assays confirmed that the expression of *pgsBCAA* was directly negatively regulated by the regulator TnrA, and the TnrA-binding site was identified in the *pgsBCAA* promoter of *B. licheniformis*. Finally, we demonstrated that the TnrA-binding sites in the *pgsBCAA* promoters were highly conserved in *Bacillus*. Taken together, our results implied that TnrA was a direct repressor for *pgsBCAA* expression in γ-PGA synthesis, and our research provided a novel regulatory mechanism underlying γ-PGA synthesis, and a new approach that deficiency of *tnrA* increases γ-PGA production.

## Materials and methods

### Bacterial strains and plasmids

The strains and plasmids used in this research were provided in **Table 1**. *B. licheniformis* WX-02 acts as the original strain for constructing mutants. The plasmid T_2_(2)-Ori was applied to construct the *tnrA* knockout vector T_2_-tnrA. The TnrA overexpression vector pHY-tnrA was obtained based on pHY300. All premiers used for strain construction were provided in **Table 2**.

**Table 1.**
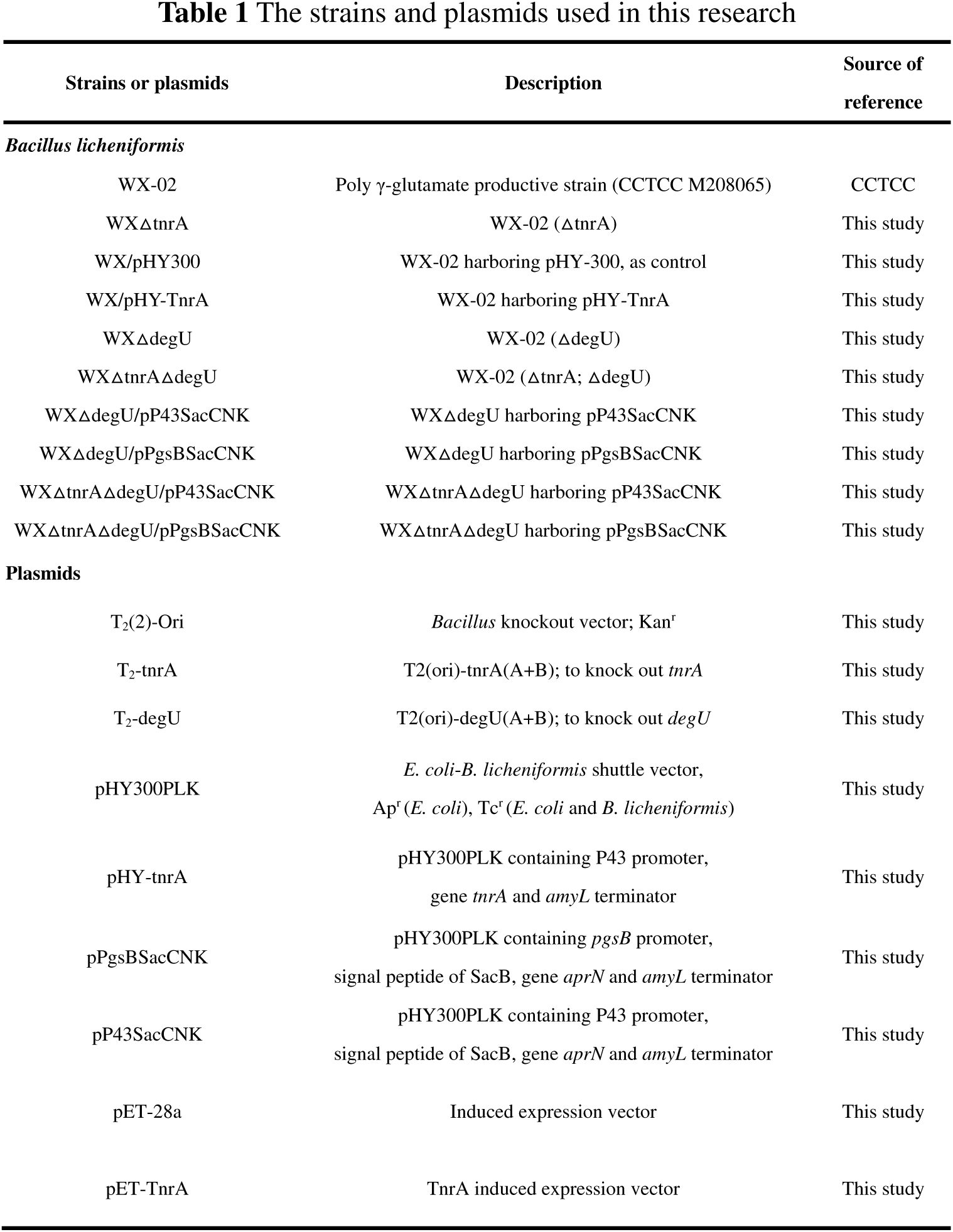
The strains and plasmids used in this research

| Strains or plasmids | Description | Source of reference |
| --- | --- | --- |
| *Bacillus licheniformis* |  |  |
| WX-02 | Poly γ-glutamate productive strain (CCTCC M208065) | CCTCC |
| WXΔtnrA | WX-02 (ΔtnrA) | This study |
| WX/pHY300 | WX-02 harboring pHY-300, as control | This study |
| WX/pHY-TnrA | WX-02 harboring pHY-TnrA | This study |
| WXΔdegU | WX-02 (ΔdegU) | This study |
| WXΔtnrAΔdegU | WX-02 (ΔtnrA; ΔdegU) | This study |
| WXΔdegU/pP43SacCNK | WXΔdegU harboring pP43SacCNK | This study |
| WXΔdegU/pPgsBSacCNK | WXΔdegU harboring pPgsBSacCNK | This study |
| WXΔtnrAΔdegU/pP43SacCNK | WXΔtnrAΔdegU harboring pP43SacCNK | This study |
| WXΔtnrAΔdegU/pPgsBSacCNK | WXΔtnrAΔdegU harboring pPgsBSacCNK | This study |
| **Plasmids** |  |  |
| T<sub>2</sub>(2)-Ori | *Bacillus* knockout vector; Kan<sup>r</sup> | This study |
| T<sub>2</sub>-tnrA | T2(ori)-tnrA(A+B); to knock out *tnrA* | This study |
| T<sub>2</sub>-degU | T2(ori)-degU(A+B); to knock out *degU* | This study |
| pHY300PLK | *E. coli*-*B. licheniformis* shuttle vector, Ap<sup>r</sup> (*E. coli*), Tc<sup>r</sup> (*E. coli* and *B. licheniformis*) | This study |
| pHY-tnrA | pHY300PLK containing P43 promoter, gene *tnrA* and *amyL* terminator | This study |
| pPgsBSacCNK | pHY300PLK containing *pgsB* promoter, signal peptide of SacB, gene *aprN* and *amyL* terminator | This study |
| pP43SacCNK | pHY300PLK containing P43 promoter, signal peptide of SacB, gene *aprN* and *amyL* terminator | This study |
| pET-28a | Induced expression vector | This study |
| pET-TnrA | TnrA induced expression vector | This study |

**Table 2.**
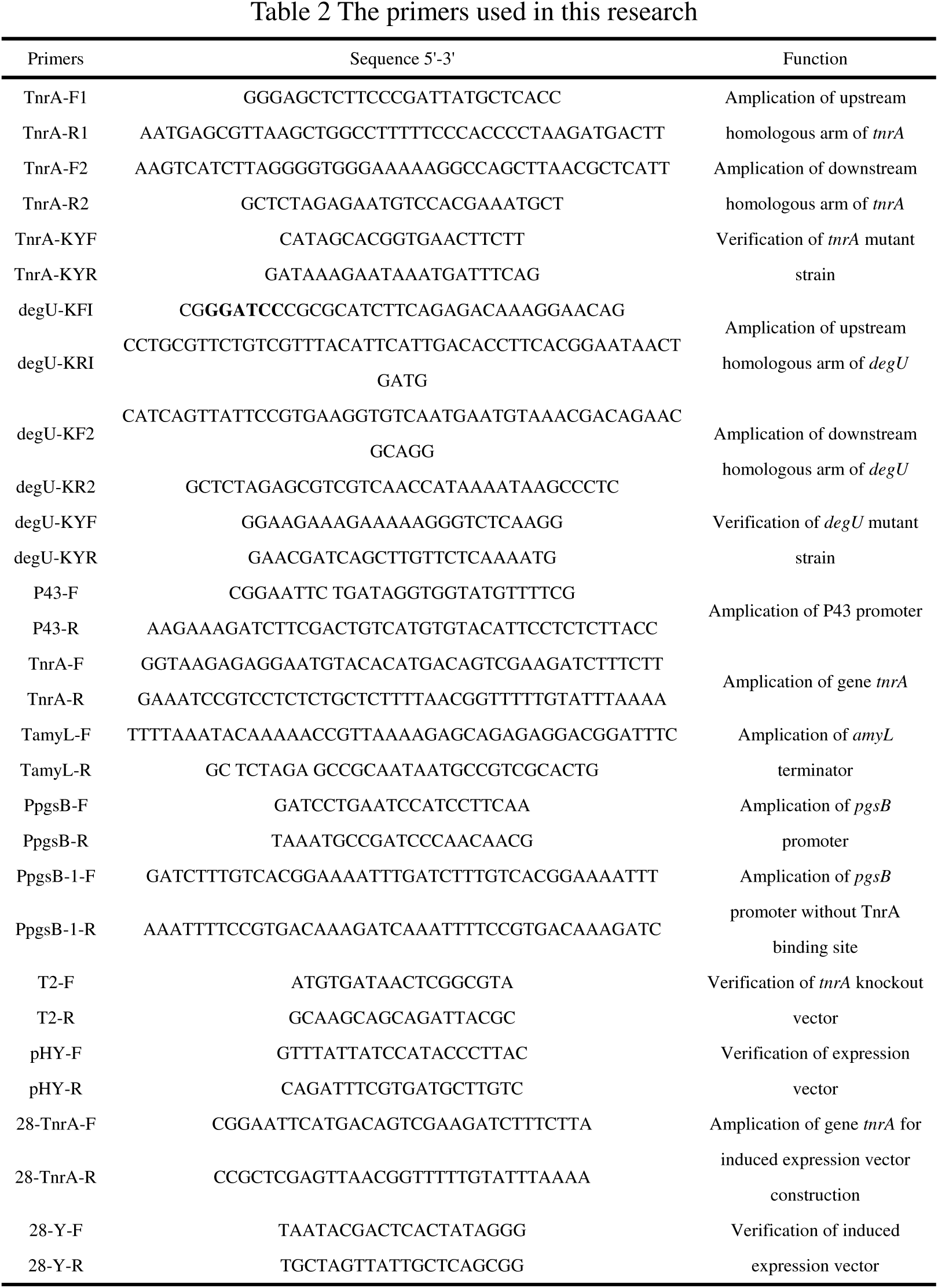
The primers used in this research

**Table 3.**
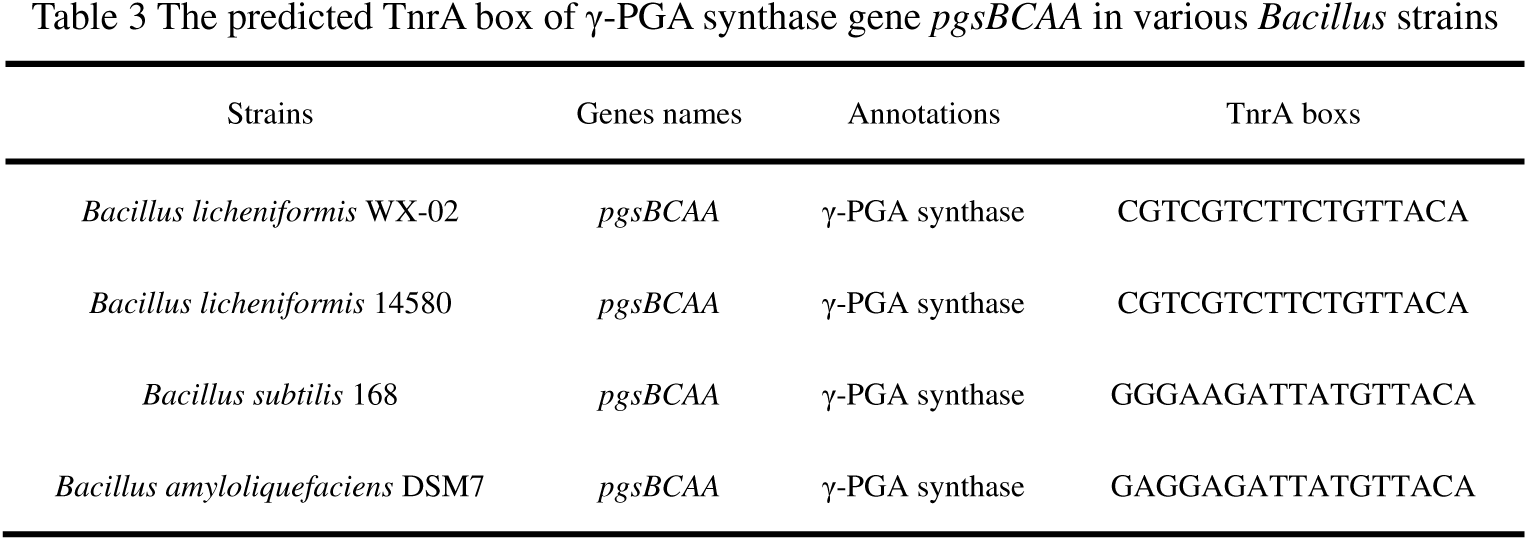
The predicted TnrA box of γ-PGA synthase gene *pgsBCAA* in various *Bacillus strains*

### Medium and cultivate condition

LB medium (10 g/L Tryptone, 5 g/L yeast extract, 10 g/L NaCl, pH 7.2) was served as the basic medium for strain cultivation, and responsible antibiotic (20 μg/mL kanamycin, 50 μg/mL ampicillin or 20 μg/mL tetracycline) was added into the medium when necessary. The seed culture of *B. licheniformis* was prepared in a 250 mL flask with 50 mL LB medium, and incubated at the rotatory shaker with 180 rpm at 37ºC for 10-12 h until OD_600_ reached 4.0~4.5, and then transferred into the γ-PGA and nattokinase production medium, respectively. The γ-PGA production medium consisted of (per liter) 80 g glucose, 30 g glutamic acid, 10 g sodium citrate, 8 g NH_4_Cl, 1 g CaCl_2_, 1 g K_2_HPO_4_⋅3H_2_O, 1 g MgSO_4_⋅7H_2_O, 1 g ZnSO_4_⋅7H_2_O and 0.15 g MnSO_4_⋅7H_2_O at pH 7.2(25). The nattokinase fermentation medium contained (per liter) 20 g glucose, 10 g peptone, 5 g yeast extract, 10 g NaCl, 10 g soy peptone, 10 g corn steep liquor, and 6 g (NH_4_)_2_SO_4_, pH 7.2 (35). The fermentation conditions were the same as those of the seed culture.

### Construction of *tnrA* deletion strain

The *tnrA* mutant strain was constructed according to our previous method (35). Briefly, the up-stream and down-stream homologous arms of *tnrA* were amplified with the corresponding premiers, and fused by Splicing-Overlapping-Extension PCR (SOE-PCR). The fused fragments was then inserted into the plasmid T_2_(2)-Ori at the restriction sites *Xba*I and *Sac*I, diagnostic PCR and DNA sequence were used to confirm that the recombinant plasmid (T_2_-tnrA) was constructed successfully.

The vector T_2_-tnrA was electro-transferred into *B. licheniformis* by the previous reported method, verified by diagnostic PCR and plasmid extraction (36). The positive transformants were cultivated in the LB medium with 20 μg/mL kanamycin at 45ºC to promote the single-cross transformation for three generations, and then transferred into the kanamycin-free medium at 37ºC for six generations. Then, the kanamycin-sensitive colonies were verified by diagnostic PCR, and the mutant strain was confirmed by DNA sequencing, named as WXΔtnrA.

### Construction of expression vectors

The expression vectors were constructed via according to the following steps, and the TnrA expression vector was served as an example. Briefly, the P43 promoter of *B. subtilis* 168 (K02174.1), gene *tnrA* (3101408) and *amyL* terminator (FJ556804.1) of *B. licheniformis* WX-02 were amplified using the corresponding primers, and fused by SOE-PCR. The fused fragments were inserted into the expression vector pHY300PLK at the restriction sites *Xba*I and *Eco*RI. Diagnostic PCR and DNA sequence were used to confirm that the recombinant plasmid (pHY-tnrA) was constructed successfully. Similarly, the nattokinase expression vector mediated by *pgsB* promoter was constructed by the same method, named as pPgsBSacCNK, and the nattokinase expression vector mediated by P43 promoter pP43SacCNK was obtained in our previous research (37).

### Analytical methods

Cell biomass was measured based on cell dry weight. The γ-PGA concentration was determined by High performance liquid chromatography (HPLC) according to the method described in our previous research (38). Glucose and glutamic acid concentrations were measured by a SBA-40C bioanalyzer according to the manual instruction (Academy of sciences, Shandong, China).

### Expression and purification of TnrA in *Escherichia coli*

The TnrA induced expression vector was constructed based on the following steps. Briefly, the fragment of *tnrA* (3101408) was amplified from *B. licheniformis* WX-02 using the corresponding premiers, and inserted into the induced vector pET-28a at the restriction sites *Eco*RI and *Xho*I. The recombinant plasmid pET-TnrA was verified by diagnostic PCR and DNA sequence. Expression and purification of TnrA were performed as described previously (39). The purified proteins were verified by sodium dodecyl sulfate-polyacrylamide gel electrophoresis (SDS-PAGE), and protein concentration was measured by a microplate reader (Bio-Rad, USA).

### Electrophoretic mobility shift assays

The sequence of *pgsBCAA* promoter (Gene ID: 3028267, from -300 to +50) for electrophoretic mobility shift assays (EMSAs) were amplified with gene-specific primers containing 5’-biotin-modified universal primer. The PCR products were analyzed by agarose gel electrophoresis and purified using a PCR purification kit (Omega, USA). The concentration of biotin-labeled DNA probe was determined using a trace Spectrophotometer NanoDrop 2000 (Thermo, USA). For amplification the *pgsBCAA* promoter without the TnrA binding site (P_*pgsB*’_), SOE-PCR was applied to fuse the upstream and downstream fragments of TnrA binding site, and formed the fragment P_*pgsB*’_.

The EMSAs were carried out according to the manufacturer’s protocol of the chemiluminescent EMSA kit (Beyotime Biotechnology, China). The binding reaction mixture containing 10 mM Tris-HCl (pH 8.0), 25 mM MgCl_2_, 50 mM NaCl, 1 mM dithiothreitol (DTT), 1 mM EDTA, 0.01% Nonidet P-40, 50 μg/mL poly(deoxyinosinic-deoxycytidylic) acid (poly(dI-dC)), and 10% glycerol. Biotin-labeled DNA probes were incubated individually with various concentrations of TnrA proteins at 25°C for 20 min. After binding, the samples were separated on a 4% nondenaturing PAGE gel in an ice-bath of 0.5×Tris-borate-EDTA (TBE) at 100 V, and trans-blotted to nylon membrane with mini trans-blot electrophoresis apparatus (Liuyi, China). Then, the membrane was treated by Chemiluminescent EMSA Kit (Beyotime, China) and analyzed with the MF-ChemiBIS (DNR Bio-imaging systems, Israel) (40).

### Transcriptional level assay

The gene transcriptional levels of mutant strains were analyzed based on the previously reported method (8). In brief, total RNA was extracted by TRIzol^®^ Reagent, and the trace DNA was digested by DNase I. The first stand of cDNA was amplified by the Revert Aid First Strand cDNA Synthesis Kit (Thermo, USA). The 16S rRNA of *B. licheniformis* WX-02 was used as the reference gene. The gene transcriptional levels of recombinant strains were compared with those of the control strain after being normalized to the reference gene 16S rRNA. All the experiments were performed in triplicates.

### Nattokinase activity assay

The nattokinase activity was measured by fibrin degradation method (35). In brief, 0.4 mL fibrinogen solution (0.72%, w/v) and 1.4 mL Tris-HCl (50 mM, pH 7.8) were mixed and incubated at 37ºC for 5 min, added 0.1 mL thrombin (20 U/mL) and incubated at 37ºC for 10 min. The prepared fibrin-substrate solution was then mixed with 0.1 mL nattokinase-containing broth, and incubated at 37ºC for 60 min and shook every 20 min during the incubation. Then, 2 mL trichloroacetic acid (TCA) solution (0.2 M) was added to stop the reaction. As a control, 0.1 mL nattokinase-containing broth and 2 mL TCA solution (0.2 M) were added into the prepared fibrin-substrate solution after incubating at 37ºC for 60 min. The samples were centrifuged at 13,000 *g* for 10 min and measured the absorbance at 275 nm. One unit nattokinase activity (FU) was defined as the amount of enzyme leading to 0.01 increase of absorbance in 1 min.

### Computational analysis

The TnrA binding sites in the *pgsBCAA* promoters of various *Bacillus* strains were identified by MEME/MAST tools (http://meme-suite.org/), according to the conserved TnrA box reported in the previous research (40).

### Statistical analyses

All samples were analyzed in triplicate, and the data were presented as the mean ± the standard deviation for each sample point. All data were conducted to analyze the variance at P < 0.05 and P < 0.01, and a *t* test was applied to compare the mean values using the software package Statistica 6.0 (35).

## Competing interests

The authors declare that they have no competing interests.

## Athour’s contribution

D Cai and S Chen designed the study. D Cai and Y Chen carried out the molecular biology studies and construction of engineering strains. D Cai, Y Chen, S Wang and F Mo carried out the fermentation studies. D Cai, X Ma and S Chen analyzed the data and wrote the manuscript. All authors read and approved the final manuscript.

## Acknowledgments

This work was supported by the National Program on Key Basic Research Project (973 Program, No. 2015CB150505), the Science and Technology Program of Wuhan (20160201010086).

## Supporting Information

All the primers sequences for RT-qPCR were listed in **Table S1**. These information were available free of charge via the Internet: http://aem.asm.org/.

